# Two morphologically distinct formae speciales in *Neonectria magnoliae* differ in their virulence on Magnolia family hosts *Liriodendron tulipifera* and *Magnolia fraseri*

**DOI:** 10.1101/2025.01.31.635987

**Authors:** Hannah M. Petronek, Shannon C. Lynch, Brian Lovett, Angie M. Martin, Danielle K.H. Martin, Matt T. Kasson

**Author notes:** Corresponding author: Matt T. Kasson. Warnell School of Forestry and Natural Resources, University of Georgia, Athens, GA 30605.

## Abstract

The family Nectriaceae includes numerous phytopathogenic fungal genera that cause canker diseases on both angiosperm and conifer hosts worldwide. Among these, *Neonectria* species are globally important canker pathogens of numerous hosts, but their roles in contributing to forest decline and mortality outside their role in beech bark disease and apple canker are largely understudied. In the U.S., *Neonectria magnoliae* causes perennial cankers on two native hosts in central Appalachia: Fraser magnolia (*Magnolia fraseri*) and tulip-poplar (*Liriodendron tulipifera*) and has been recently confirmed from non-native star magnolia (*Magnolia stellata*) in West Virginia. Both native hosts occur in the central Appalachian Mountains, but Fraser magnolia occurs mostly at higher elevations. *Neonectria magnoliae* was first described in 1943, yet its impact across the forested landscape remains unclear. To clarify host-specific differences across the contemporary range of *Neonectria magnoliae*, we used multi-locus phylogenetics, comparative pathogenicity / virulence assays, and morphological analyses to determine if *N. magnoliae* represents two cryptic species that specialize on tulip-poplar and magnolia, or if *N. magnoliae* has host-specific pathotypes. Our studies revealed two morphologically distinct formae speciales within *N. magnoliae*: 1) *Neonectria magnoliae* f. sp. *liriodendri;* strains originating from tulip-poplar with increased virulence on this host and lacking macroconidia production and 2) *Neonectria magnoliae* f. sp. *magnoliae;* strains originating from Fraser magnolia with increased virulence on this host and producing macroconidia readily in culture. Overall, the incidence of these two pathotypes indicates that neither pathogen alone poses serious risks to either host but adds to cumulative stresses that both tree species are experiencing in the face of global climate change.

## INTRODUCTION

The Nectriaceae is a hyper-diverse fungal family containing >1,300 species across >70 genera with diverse ecologies including mycoparasites, entomopathogens, and phytopathogens (Lombard et al. 2015; Stauder et al. 2020). Among the phytopathogenic Nectriaceae, *Calonectria, Fusarium, Neocosmospora,* and *Neonectria* genera comprise nearly half of the described taxa for the family and contain a majority of the species that are known to cause stem canker diseases of trees. *Neonectria* contains more than 20 species, a majority of which cause annual and perennial canker diseases on diverse angiosperms and gymnosperms around the globe (Chaverri et al. 2011; Petronek et al. 2024). *Neonectria* have asexual states in *Cylindrocarpon* Wollenw. (Brayford et al. 2004; Mantiri et al. 2001; Rossman et al. 1999) and their sexual state is characterized by bright red, globose perithecia that often appear in clusters on bark tissues and exposed wood.

Many of these perithecia-producing *Neonectria* fungi cause significant cankers on hardwood hosts. In North America, *Neonectria ditissima* and *Neonectria faginata* are among the most well-known species, as both are causal agents of beech bark disease of American beech (*Fagus grandifolia* Ehrh.) (Cale et al. 2017). *Neonectria ditissima* causes perennial target cankers on non-beech hosts, encompassing the genera *Acer, Betula, Ilex, Juglans, Sorbus, Populus,* and *Salix* among others (Stauder et al. 2020). In contrast, *N. faginata* is restricted to American beech following colonization by the beech scale insect (*Cryptococcus fagisuga* Baer.) (Morrison et al. 2021). In addition to exhibiting a narrow host range, *N. faginata* has never been found outside the U.S. or decoupled from beech bark disease (Cale et al. 2017; Morrison et al. 2021; Mahoney et al. 1999). In addition to these two species, *N. magnoliae* and *N. punicea*, as well as the unrelated *Cosmospora obscura* and *Thelonectria veuillotiana –* two other red perithecia producing Nectriaceae species *–* have been confirmed on tree species from these same forests and same hosts (Stauder et al. 2020), which has led to confusion as to which fungus was ultimately the causal agent of disease (Booth 1959; Castlebury et al. 2006). Prior to DNA barcoding, identifications were made primarily based on perithecial characteristics, ascospore size, and micro- and macroconidia measurements for the *Cylindrocarpon* state (Booth 1959; Lohman & Watson 1943; Castlebury et al. 2006). However, some species have overlapping measurements and as such their identification cannot be fully resolved based on morphological characteristics alone.

In the 1930s, Lohman & Watson (1943) discovered a ‘Nectria’ canker pathogen of tulip-poplar (*Liriodendron tulipifera* L.) (FIG. 1A-B), Fraser magnolia (*Magnolia fraseri* Walter) (FIG. 1D-E), and umbrella magnolia (*Magnolia tripetala* (L.) L.) that they described as *Nectria magnoliae.* It was found to be morphologically distinct from *N. ditissima* and previously thought to be intermediate between *N. ditissima* and *N. coccinea,* the causal agent of beech bark disease on European beech (*Fagus sylvatica* L.). In 2006, Castlebury and colleagues synonymized *Nectria magnoliae* with *Neonectria ditissima* based on a single strain (CBS 118919) from tulip-poplar in Tennessee but did not include *N. magnoliae* strain CBS 380.50 from tulip-poplar in North Carolina. Stauder et al. (2020) conducted a wider survey of tulip-poplar and Fraser magnolia across West Virginia, recovering numerous isolates (FIG. 1D,H) from several locations. *N. magnoliae* formed a well-supported monophyletic clade within *Neonectria,* separate from *N. ditissima* and strain CBS 118919. An independent study by Grafenhan et al. (2011) also supported these findings: *N. magnoliae* strain CBS 380.50 from tulip-poplar in North Carolina grouped separately from *N. ditissima*. Taken together, these findings indicate that both *N. magnoliae* and *N. ditissima* are canker pathogens of tulip-poplar. Yet, the range of both species on tulip-poplar, especially *N. ditissima*, remains unclear due to few collections and sequence data.

**Figure 1:**
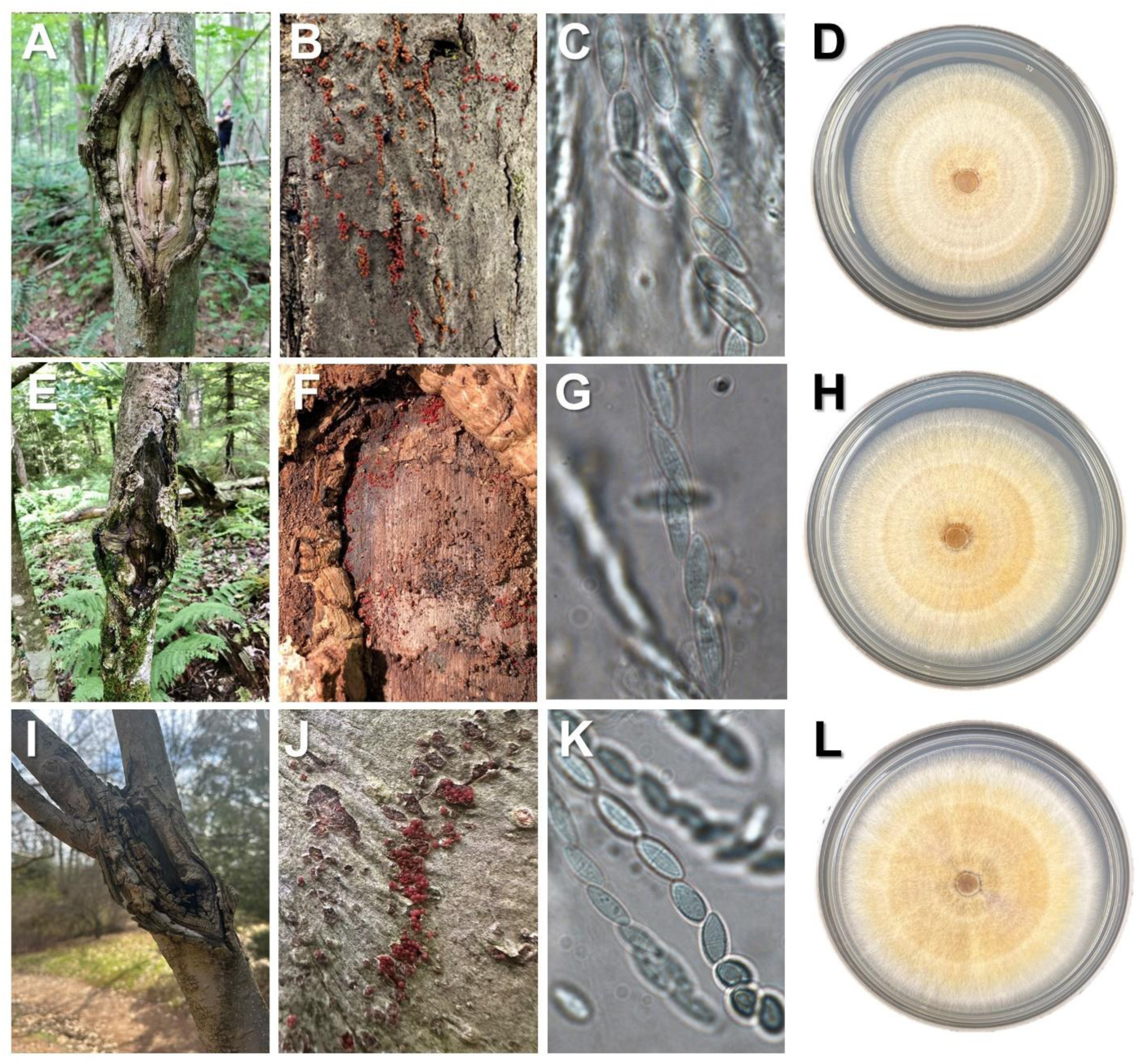
*Neonectria magnoliae* cankers (A,E,I), sexual fruiting bodies or perithecia (B,F,J), asci with uniseptate ascospores (C,G,K) and cultures of the C*ylindrocarpon* anamorph (D,H,L) derived from single ascospore colonies. Hosts include tulip-poplar (*Liriodendron tulipifera*; A-D), Fraser magnolia (*Magnolia fraseri*; E-H) and star magnolia (*Magnolia stellata*; I-L) sampled across West Virginia, USA. Culture age ∼2-3 wks on PDA.

Spaulding et al. (1936) conducted cross-inoculation trials on tulip-poplar, sugar maple, red maple, sweet birch, swamp birch, American beech, black walnut, bigtooth aspen, white oak, black oak, and pignut hickory in New Hampshire and North Carolina to investigate the pathogenicity of *N. magnoliae, N. ditissima,* and other ‘Nectria’ species. Of the hosts tested between the two study locations, *N. magnoliae* caused cankers on tulip-poplar trees only, and yielded significantly smaller cankers compared to multiple strains of *N. ditissima* on this host (Lohman & Watson 1943). Stauder et al. (2020) included *N. magnoliae* strains from tulip-poplar in field inoculation studies to compare pathogenicity and virulence of *N. faginata, N. ditissima*, and *N. magnoliae* on various non-beech hosts, including tulip-poplar, but not magnolia species. While select *N. ditissima* strains colonized and caused minimal cankers on tulip-poplar, the cankers produced by *N. magnoliae* were significantly larger compared to *N. ditissima* and *N. faginata*. The study by Stauder and colleagues was the first to confirm pathogenicity and virulence of molecularly confirmed strains of *N. magnoliae.* Given that both *N. ditissima* and *N. magnoliae* are confirmed canker pathogens of tulip-poplar, it remains unclear if isolates used by Spaulding et al. (1936) were the former or the latter, and if *N. ditissima*, were the strain(s) used isolated from tulip-poplar.

Much of the previous work on *N. magnoliae* has focused on interactions with tulip-poplar on account of its economic importance as a forest species as well as its large geographic distribution across the eastern U.S. Recently, a scaffold-level genome and assembly for *N. magnoliae* strain NRRL 64651 from this host in West Viriginia was published to support ongoing work on this canker pathogen (Petronek et al. 2024). Stauder et al. (2020) uncovered morphological and genetic differences between strains originating from tulip-poplar and Fraser magnolia. However, limited availability of strains of *N. magnoliae* from Fraser magnolia compared to tulip-poplar strains prevented in-depth comparisons at the time (Stauder et al. 2020). Regardless, differences in conidia and ascospore morphology between strains from either host were noted. For instance, macroconidia were never observed for tulip-poplar strains but readily developed in Fraser magnolia strains of *N. magnoliae,* while microconidia readily formed in strains originating from both hosts. Interestingly, Lohman & Watson (1943) reported that some *N. magnoliae* cultures failed to produce macroconidia, but did not specify host origin for these strains. Additionally, Stauder et al. (2020) observed that the target-shape pattern of cankers characteristic of this genus was less pronounced on Fraser magnolia stems (FIG. 1E). Fraser magnolia trees grow at elevations ranging from 600 meters to 1,700 meters (Delia-Bianca 1965), while tulip-poplar trees grow at elevations < 300 meters up to 1,370 meters (Beck 1962). As both species belong to the family Magnoliaceae and exhibit both geographical and elevational overlap, further investigation of the distribution of *N. magnoliae* across both hosts was warranted.

In this study, we sought to clarify host-specific differences across the contemporary range of *N. magnoliae* in West Virginia and determine if *N. magnoliae* represents two cryptic species that specialize on tulip-poplar or Fraser magnolia, or if *N. magnoliae* has host-specific pathotypes. To accomplish this, we sought to: 1) bolster existing culture collections of *N. magnoliae* from both tulip-poplar and Fraser magnolia with sampling efforts focused across West Virginia; 2) phylogenetically resolve species boundaries for a comprehensive set of *N. magnoliae* strains; 3) conduct morphological and cross-pathogenicity studies of *N. magnoliae* originating from both hosts in West Virginia.

## MATERIALS AND METHODS

### Field collections of N. magnoliae

Sampling locations were selected based primarily on previous survey findings reported by Stauder et al. (2020) as well as other historic observations. *N. magnoliae* was sampled from 7 counties in West Virginia, with numerous tulip-poplar isolates sampled from Snake Hill Wildlife Management Area in Monongalia County and many Fraser magnolia isolates from Gaudineer Knob National Recreation Area in Randolph County (TABLE 1). For each host, samples were taken and isolates obtained from 2+ counties.

**Table 1.**
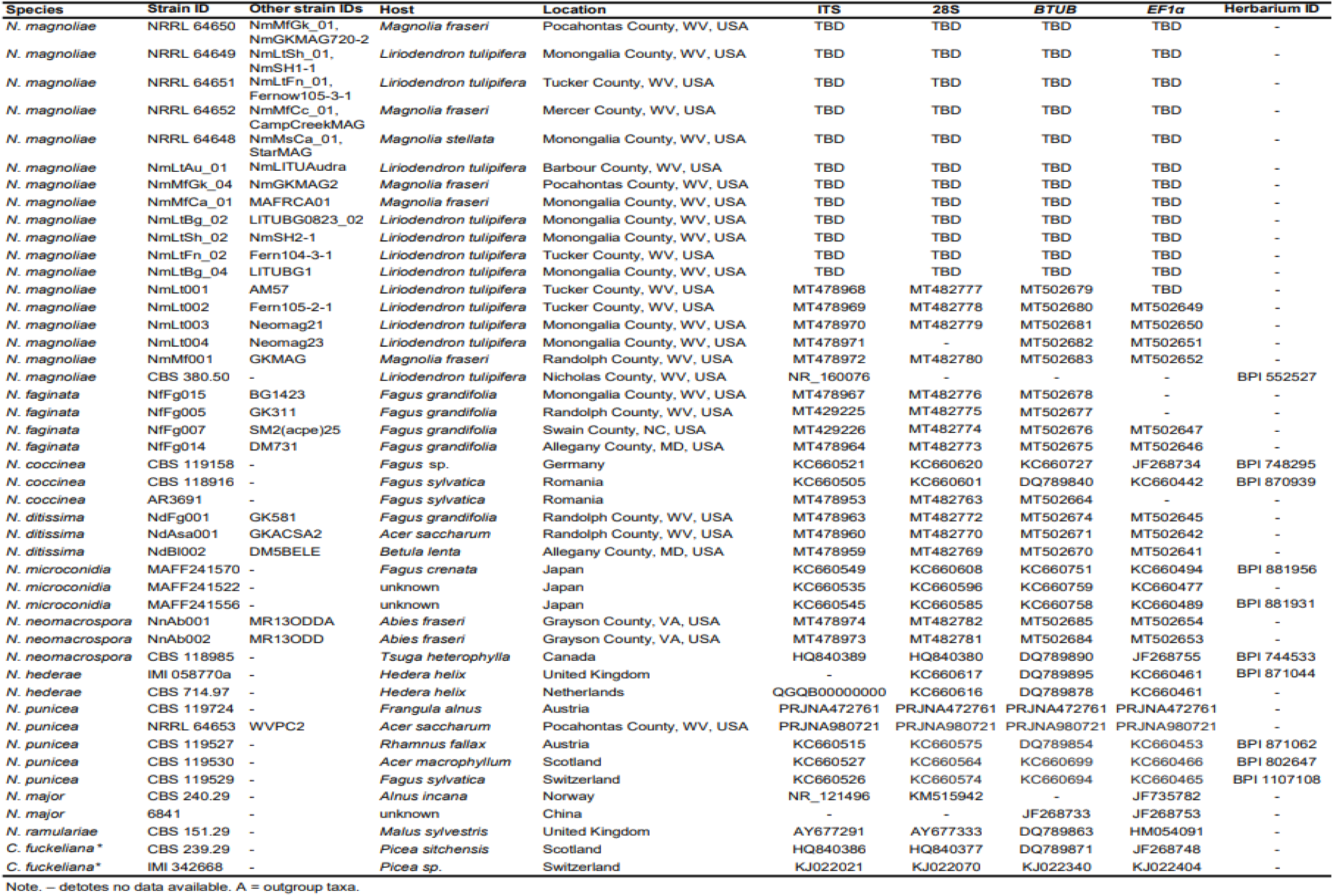
Species and isolates used in the single-locus and multi-locus phylogenetic analysis and their associated metadata.

### Isolation of N. magnoliae

From 1-cm diameter bark discs, 3-5 perithecia were removed with the use of a sterile scalpel or teasing needle and squashed in a 1.5 mL microcentrifuge tube containing 1 mL of sterile, distilled water with a micropestle. After maceration, microcentrifuge tubes were vortexed for 15 s and then 100 µL of the spore suspension spread with a sterile cell spreader on potato dextrose agar (PDA) plates amended with streptomycin sulfate (10 mg/1000 mL) and tetracycline hydrochloride (100 mg/1000 mL) antibiotics. Plates were checked after 24 and 48 hours for germinating ascospores. Five germinated ascospores per plate were sub-cultured onto new PDA plates, and one isolate was selected for long-term use and storage.

### DNA extraction & molecular characterization

DNA was extracted from 13 representative strains of *Neonectria magnoliae* collected across 8 sampling locations in West Virginia (TABLE 1) according to the protocol described in Stauder et al. (2020). To confirm species identities for all *N. magnoliae* strains and evaluate inter- and intraspecies relatedness, we generated Sanger DNA sequences for each of the following loci: internal transcriber spacer (*ITS*), large subunit rRNA (*28S*), translation elongation factor 1-alpha (*EF1-a*), and beta-tubulin (*BTUB*). Primer names, sequences, and PCR thermocycling conditions are listed in SUPPLEMENTARY TABLE 1. PCR reagents and post-PCR cleanup are as described in Lovett et al. (2024). Sanger sequences were generated by Eurofins Genomics (Louisville, KY, USA) using the same primers in our PCR reactions.

To determine if *N. magnoliae* comprised two or more distinct species, phylogenetic analyses were conducted across 18 *N. magnoliae* isolates from two known hosts across multiple geographic locations, including the type specimen for *N. magnoliae* (CBS 380.50; Lohman & Watson 1943). Isolates from a previously unknown non-native host, star magnolia (*Magnolia stellata* Maxim.) (FIG. 1L). DNA sequence data for representative *Neonectria* species were pulled from NCBI GenBank as described in TABLE 1. Included sequence data and associated accession numbers are listed in TABLE 1. Nucleotide sequences for *N. punicea* NRRL 64653 *ITS, 28S, EF1-ɑ*, and *BTUB* were extracted from our recently assembled genome (Petronek et al. 2024) using *Neonectria* reference sequences for *ITS* (MT478967), *28S* (MT482776), *BTUB* (MT502678), and *EF1-ɑ* (MT502647). Datasets for each of the 4 individual loci and combined (2-locus and 4-locus) datasets were analyzed using maximum likelihood & Bayesian inference for a total of 12 analyses.

Chromatograms were quality-checked using default parameters and clipped in Geneious Prime ® 2023.1.2 (http://www.geneious.com/). Each locus was aligned separately using MAFFT on the GUIDANCE2 server (https://taux.evolseq.net/guidance/); Penn et al. 2010; Sela et al. 2015). Residues with GUIDANCE confidence scores <0.5 were masked with an N. Nucleotide substitution models were selected in MEGA 11.0.13 using the ‘Model Test’ feature and the subsequent Akaike information criterion (AICc) scores (Kumar et al. 2016). The determined optimal model was HKY for all single-gene trees and both concatenated trees. Individual gene alignments for *EF1-ɑ, BTUB1, ITS*, and *LSU/28S* were performed. Then, these 4 alignments were used in maximum likelihood (ML) analyses (RAxML 8.2.12; Stamakis 2014) with 1000 bootstraps and Bayesian inference (BI) analyses (MrBayes 3.2.5; Ronquist et al. 2012), using default parameters, unless otherwise noted (Stajich et al. 2024; also see code and notes in Data Availability Statement. Trees were viewed and prepared for publication using Geneious Prime, FigTree v1.3 (http://tree.bio.ed.ac.uk/software/figtree/), and Inkscape 1.3.2 (https://www.inkscape.org/). The ML two-locus and four-locus phylogenetic trees were annotated with BI support values where topology was in agreement.

### Tulip-poplar pathogenicity / virulence assay

Field inoculations were conducted in October 2020 to investigate the pathogenicity and virulence of *Neonectria magnoliae* strains originating from tulip-poplar and Fraser magnolia on tulip-poplar at West Virginia University’s Research Forest near Bruceton Mills, WV. This site was selected due to its accessibility and density of young, tulip-poplar trees. Four *N. magnoliae* strains originating in West Virginia were selected for inoculation: strains NRRL 64649 and NRRL 64651 from tulip-poplar and strains NRRL 64650 and NmGKMAG5 from Fraser magnolia. All study isolates were grown in pure culture on PDA for 2-3 weeks at room temperature (FIG. 1D,H,L). Prior to inoculations, a sterile 1-cm cork borer was used to cut fully colonized inoculation plugs. Negative control plugs were cut from sterile PDA plates.

A total of 25 tulip-poplar trees were selected and flagged for inoculations, ensuring obvious cankers, significant external stem defects, or dieback were not present prior to inoculation. Diameter at breast height (DBH) was measured for each tree. Each strain was inoculated into 5 trees in triplicate (3 inoculations per tree) for a total of 15 inoculation sites per *N. magnoliae* strain. Negative control trees received sterile agar plugs. Inoculations were made by using a 1-cm diameter sterile leather punch and hammer to excise bark plugs, replacing the plug with colonized agar plugs, and securing plugs with 1” lab tape. Each tree was labeled and inoculated at 50, 100, and 150 cm from the soil line.

At 6 months post-inoculation (MPI), trees were destructively sampled by removing the bark around the inoculation site with a chisel. Canker length and width were measured to permit canker area calculations. Additional data were recorded, including overall tree health and presence of perithecia. To fulfill Koch’s postulates, one canker per tree was microsampled 4 times with a flame-sterilized bone marrow biopsy tool. Plugs were kept cool until brought back to the laboratory where they were surface sterilized in 10% commercial bleach and plated on PDA. The presence of fungal growth was recorded and subcultured as it was observed. Efforts to re-isolate fungi were abandoned after 14 days. Representative colonies were examined morphologically to confirm *N. magnoliae*.

### Fraser magnolia pathogenicity / virulence assay

To complement the tulip-poplar field assay, field inoculations were conducted in October 2023 to further investigate pathogenicity and virulence of *N. magnoliae* strains from tulip-poplar and Fraser magnolia on Fraser magnolia in the Monongahela National Forest adjacent to the Gaudineer Knob Recreation area on the western side of FR27. This work was done with permission from the USDA Forest Service under permit no. 008809. The Fraser magnolia field site was selected due to its accessibility, high density of host plants, and natural occurrence of *N. magnoliae* in the area. The same four *N. magnoliae* isolates used in the tulip-poplar pathogenicity / virulence assay were used. Cultures were grown and prepared for inoculations as in the tulip-poplar assay. A total of 25 Fraser magnolia trees were inoculated as described above, and the experiment was taken down 6 months post-inoculation as described above. Similar to the tulip-poplar assay, the experiment spanned the host’s dormant season, a time when perennial target canker pathogens are most active.

### Statistical analysis of canker area

A one-way ANOVA was performed to test the effect of different strains of *N. magnoliae* on canker area on Fraser magnolia trees and tulip-poplar trees using the function lm() from the “lmtest” package (Zeileis & Hothorn 2002) in R (version 2023.03.1, R Core Team 2018), using RStudio. Due to violation of ANOVA assumptions, we used the “MASS’’ package to perform a boxcox transformation (Venables & Ripley 2002). After transformation, we used Tukey’s HSD test to compare treatment means in the package “agricolae” using the HSD.test() function (Mendiburu & Yaseen 2020).

### Morphological characterization of N. magnoliae from diverse hosts

Single ascospore-derived colonies of *N. magnoliae* were grown out on PDA plates at room-temperature for approximately 4-8 weeks to allow time for sporodochial development. Microconidia (n=50) associated with the the *Cyclindroncarpon*-like anamorph were measured for nine isolates listed in SUPPLEMENTARY TABLE 2, including four isolates from Fraser magnolia, three isolates from tulip-poplar, and two isolates from star magnolia (FIG. 1D,H,L). Sporodochial masses were transferred with a sterile teasing needle and mounted onto slides prepared with a lactic acid+cotton blue mountant. Length and width measurements were recorded for 50 microconidia per isolate. All measurements were taken with a Nikon Eclipse E600 compound microscope (Nikon Instruments, Melville, NY, USA) equipped with a Nikon Digital Sight DS-Ri1 microscope camera and Nikon NIS-Elements BR3.2 imaging software.

To measure ascospores, mature perithecia were processed from field-sampled bark plugs including two perithecia from each of three hosts (SUPPLEMENTARY TABLE 3). This included tulip-poplar, Fraser magnolia, and star magnolia, a previously unreported non-native host for *N. magnoliae* (FIG. 1C,G,K). Singleton to small clusters of adjacent perithecia were excised from the colonized bark plugs using a sterile scalpel and squash-mounted on glass microscope slides prepared with a lactic acid+cotton blue mountant and sealed with clear nail polish. The length of 25 ascospores was recorded for each of the 6 slides, using the same equipment as referenced above.

### Statistical analyses of spore measurements

For ascospore measurements, a one-way ANOVA was performed to examine the differences in ascospore length grouped by the host species for the strains of *N. magnoliae* (tulip-poplar, Fraser magnolia, or star magnolia). ANOVA assumptions were assessed using the leveneTest() function in the package “car” and the shapiro.test() function in the base R “stats” package (Fox & Weisberg 2019). A Tukey’s HSD test was performed to compare treatment means in the package “agricolae” using the HSD.test() function (Mendiburu & Yaseen 2020). To correct for multiple comparisons, a Bonferroni correction was performed using the LSD.test() function in the package “agricolae”.

For microconidia measurements, a one-way ANOVA was performed to examine the differences in microconidia length and width grouped by the host species for the included strains of *N. magnoliae*. For microconidia width measurements, a one-way ANOVA was conducted using the lm() function from the “lmtest” package. ANOVA assumptions were tested using the “DHARMa” package prior to conducting a Tukey’s HSD test to compare treatment means (Hartig 2022). To correct for multiple comparisons, we conducted a Bonferroni correction. For microconidia length measurements, a 1-way ANOVA was conducted. Upon conducting tests for homogeneity of variance and normality of residuals, it was found that neither of these assumptions were met. Therefore, a square root transformation was applied to the microconidia length measurements. Despite the square root transformation, unequal variances were still noted. To remedy this, we used the Games-Howell test–a robust post-hoc test sensitive to unequal variances–using the function games_howell_test() in the package “rstatix” (Kassambara 2023).

## RESULTS

### Field surveys of N. magnoliae

*N. magnoliae* was sampled from 7 counties in West Virginia including Barbour (on tulip-poplar) and Mercer (on Fraser magnolia) Counties, which were not previously surveyed for this fungus (Stauder et al. 2020). *Neonectria magnoliae* was also recovered from a symptomatic star magnolia (*Magnolia stellata*) in West Virginia University’s Core Arboretum in Morgantown, West Virginia. This is the first report of *N. magnoliae* on any non-native magnolia species in the U.S. and the first report of this fungus on any *Magnolia* sp. outside the Allegheny Mountains region of the Appalachian Plateau.

### Phylogenetic analysis of the genus Neonectria

Both methods of phylogenetic inference -ML and BI-resolved a majority of *Neonectria* species as strongly supported (>99% bootstrap support / >0.98 posterior probability) monophyletic lineages for both the single-locus (*EF1-α, BTUB*) and the concatenated 2-locus (*EF1-α + BTUB*) and 4-locus (*EF1-α + BTUB* + *ITS +28S*) analyses with the exception of *N. punicea* and *N. major* (FIG. 2-3, SUPPLEMENTARY FIG. 1A-B). Both *ITS* and *28S* single-locus analyses were unable to resolve most of the included *Neonectria* species, with the exception of *N. neomacrospora* and *N. magnoliae*. *ITS* (>92 / >0.99) provided better support for *N. magnoliae* compared to *28S* (>74 / >0.89) (SUPPLEMENTARY FIG. 1C-D). Only the *ITS* dataset included the type strain of *N. magnoliae* (CBS 380.50) from tulip-poplar, which fell within the well-supported *N. magnoliae* clade (FIG. 3, SUPPLEMENTARY FIG. 1C).

**Figure 2.**
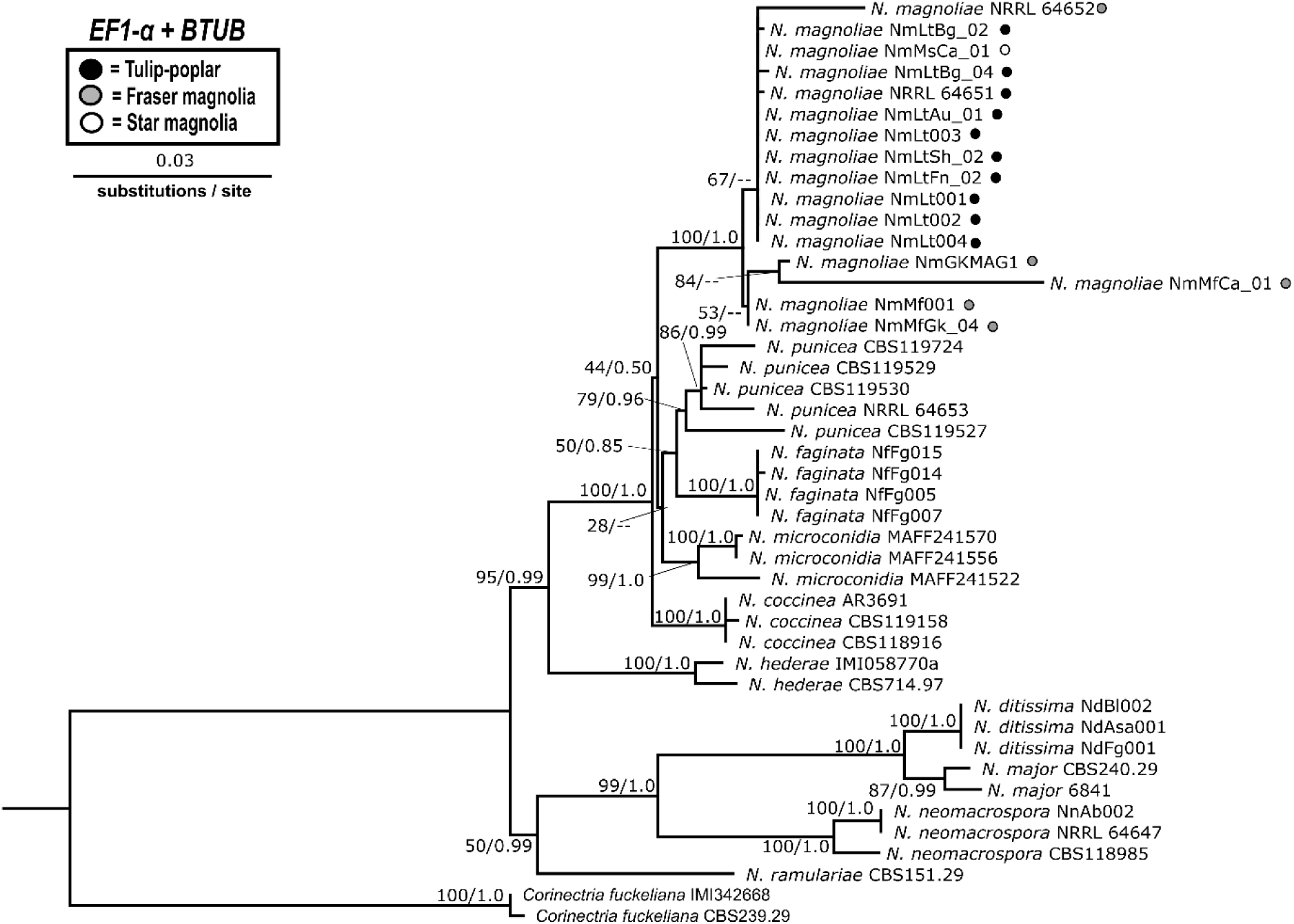
Two-locus (*EF1-α, BTUB*) concatenated phylogeny of *Neonectria* spp. with *Corinectria fuckeliana* outgroup. Topology and branch lengths shown are from the ML analysis. For each node supported in the ML analysis, bootstrap support and posterior probabilities are indicated (ML/BI) with one exception. Bootstrap support values < 50 for ML analysis are not shown nor are their corresponding BI posterior probability values. Dashes indicate that the node did not appear in the BI analysis. Strain metadata including GenBank accession numbers are listed in TABLE 1. Host origin is shown for each *N. magnoliae* strain using filled circles.

**Figure 3.**
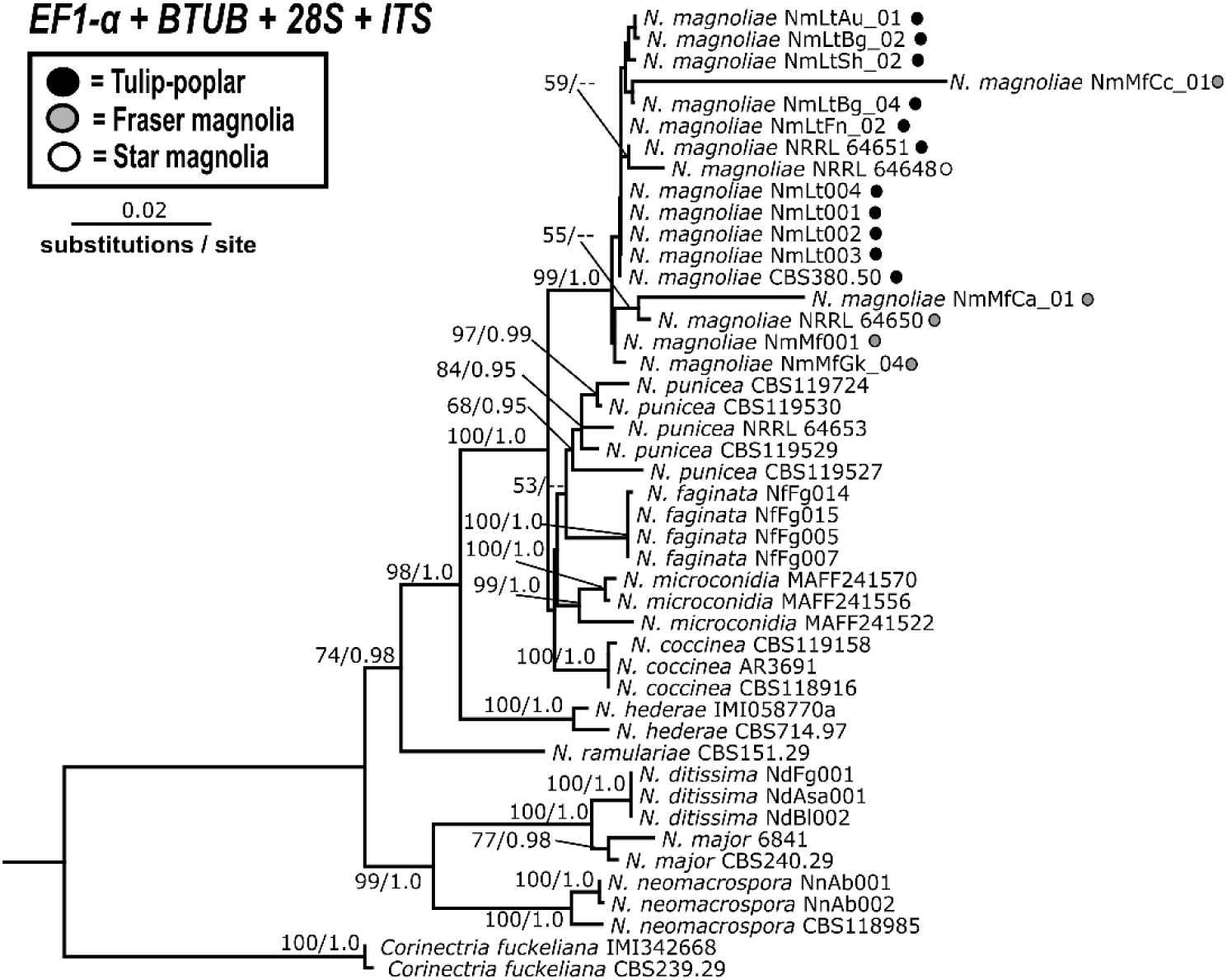
Four-gene (*ITS, LSU, EF1-α, BTUB*) concatenated phylogeny of *Neonectria* spp. with *Corinectria fuckeliana* outgroup. Topology and branch lengths are from the ML analysis. For each node supported in the ML analysis, bootstrap support and posterior probabilities are indicated (ML/BI) with one exception. Bootstrap support values < 50 for ML analysis are not shown nor are their corresponding BI posterior probability values. Dashes indicate that the node did not appear in the indicated analysis. Strain metadata including GenBank accession numbers are listed in TABLE 1. Host origin is shown for each *N. magnoliae* strain using filled circles.

The *EF1-α* and *BTUB* single-locus analyses (SUPPLEMENTARY FIG. 1A-B) and their concatenated 2-gene analysis (FIG. 2) further resolved some intraspecies differences in *N. magnoliae,* forming either host (tulip-poplar or magnolia) specific lineages or single-host dominated lineages including: 1) a moderately-supported (66%/--) tulip-poplar dominated lineage in the *EF1-α* (SUPPLEMENTARY FIG. 1A) and *EF1-α + BTUB* (FIG. 2) trees containing all tulip-poplar strains along with two magnolia-origin strains, and 2) a moderately-supported (67%/--) Fraser magnolia specific lineage in the *BTUB* (SUPPLEMENTARY FIG. 1B) tree containing three isolates from a single location in Randolph Co., WV. Bayesian analysis did not provide support for these intraspecies groupings.

Three *N. magnoliae* isolates including NRRL 64652 (Fraser magnolia, Mercer Co., WV), NmMfCa_01 (Fraser magnolia, Monongalia Co., WV), and NRRL 64648 (star magnolia, Monongalia Co., WV) appeared on unusually long branches in single locus and / or multi-locus trees including: *28S* (NRRL 64652 and NRRL 64648), *EF1-α* (NmMfCa_01), *BTUB* (NRRL 64652), *EF1-α+BTUB* (NRRL 64652 and NmMfCa_01), and *ITS+28S+EF1-α+BTUB* (NRRL 64652 and NmMfCa_01). These three magnolia-origin strains are the same strains that fell within with the tulip-poplar clade.

### Comparative pathogenicity / virulence experiments & exploration of host range

Because Lohman & Watson (1943) and Stauder et al. (2020) conducted pathogenicity tests and virulence assays using single host isolates only against tulip-poplar, our field inoculations sought to duplicate these previous efforts while also testing additional hosts including Fraser magnolia with strains of *N. magnoliae* from both hosts.

Following removal of bark from inoculated tulip-poplar trees, evidence of both necrosis and associated vascular streaking was observed (FIG. 4A-C). However, only areas of necrosis were measured, as streaking was a host-specific response to wounding as evidenced by the occurrence in all treatments including negative controls, which lacked necrosis. Streaking was completely absent from Fraser magnolia trees (FIG. 4D-F).

**Figure 4.**
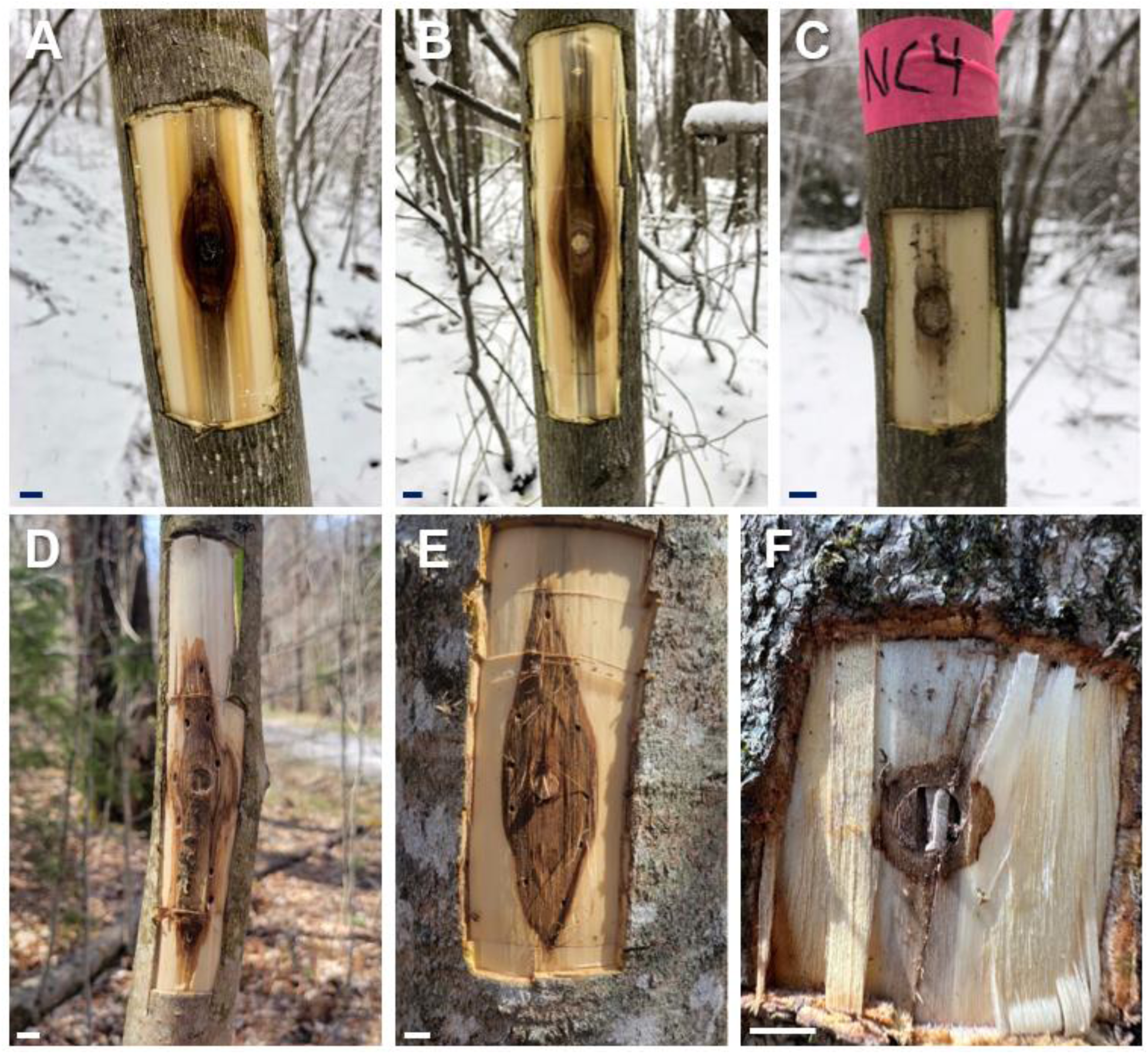
Representative *N. magnoliae* cankers from pathogenicity assays on tulip-poplar (A-C) and Fraser magnolia (D-F), 6 months post inoculation. Living stems of both tree species were inoculated with strains from tulip-poplar (A, NRRL 64649 and D, NRRL 64651) or Fraser magnolia (B, NRRL 64650 and E, NmGKMAG5), or were negative controls inoculated with a sterile PDA agar plug (C and F). Scale bars = 1 cm.

When inoculated in tulip-poplar, all *N. magnoliae* treatments caused cankers that were significantly larger than the negative controls, regardless of host-origin or strain (FIG. 5A). A significant difference in canker area was observed between isolates originating from either tulip-poplar or Fraser magnolia hosts (*F*_2,71_ = 71.79, *P* < 0.0001) but not between strains from a single host (data not shown). Mean canker area for stems inoculated with *N. magnoliae* strains from tulip-poplar was 201.80 cm^2^ and 144.42 cm^2^ for strains from Fraser magnolia (FIG. 5A). Negative control trees had a mean canker area of 22.92 cm^2^.

**Figure 5.**
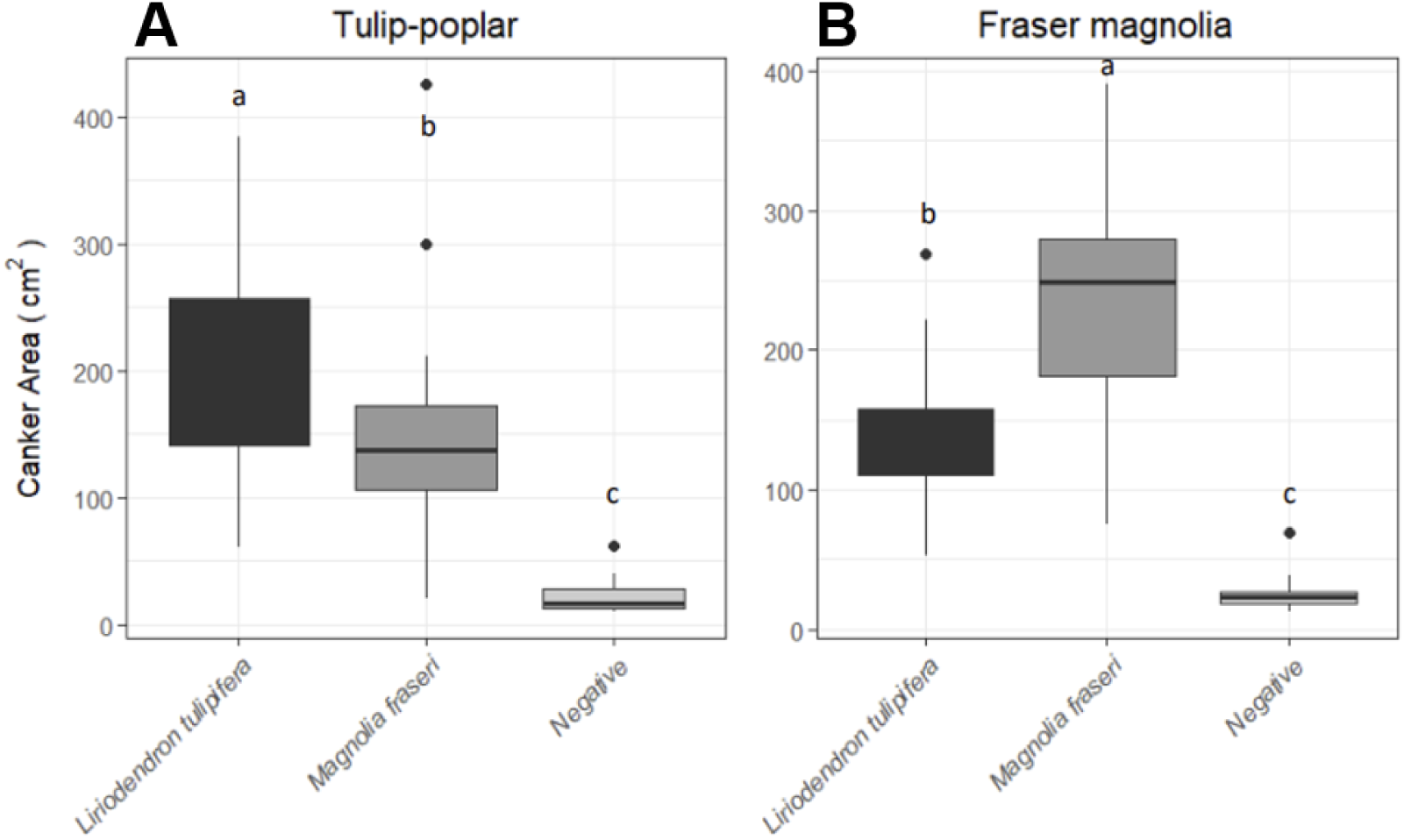
Boxplots of canker area (cm^2^) resulting from artificial inoculations of *N. magnoliae* on live tulip-poplar (A) and Fraser magnolia (B). Strains are grouped by host origin (X-axes). Letters above plots indicate significant differences.

When inoculated in Fraser magnolia, all *N. magnoliae* treatments regardless of host origin caused cankers that were significantly larger than the negative controls (FIG. 5B). Similar to the tulip-poplar pathogenicity study, significant differences in canker area were observed between isolates originating from either tulip-poplar or Fraser magnolia hosts (*F*_2,72_ = 165.89, *P* < 0.0001) but not between strains from a single host (data not shown). Mean canker area for stems inoculated with *N. magnoliae* strains originating from Fraser magnolia host was 242.27 cm^2^ and 135.94 cm^2^ for strains originating from tulip-poplar (FIG. 5B). Negative control trees had a mean canker area of 25.59 cm^2^.

In both experiments, representative microsampled canker tissues taken from one randomly chosen canker for every tree across all treatments were used to reisolate *N. magnoliae* in order to fulfill Koch’s postulates. Examination of *N. magnoliae* isolates recovered from these plugs confirmed each pathotype was associated with the correct treatments: only *N. magnoliae* strains originating from magnolia produced macroconidia in culture. Isolations from streaking only from negative control tulip-poplar never yielded *Neonectria* sp.

Based on the results reported here, two distinct pathotypes are reported: 1) strains originating from tulip-poplar with increased virulence on this host, but also pathogenic on Fraser magnolia and 2) strains originating from Fraser magnolia with increased virulence on this host, but also pathogenic on tulip-poplar. Isolates originating from star magnolia could not be delimited as comparative pathogenicity and virulence assays were not undertaken as part of this study due to the later discovery of this novel host association.

### Morphological characterization of N. magnoliae

Morphological studies were conducted to provide a comprehensive comparison of the two putative host-specific pathotypes conducted across multiple *N. magnoliae* isolates from 3 known hosts (tulip-poplar, Fraser magnolia, and star magnolia) across West Virginia (SUPPLEMENTARY TABLE 2-3). Microconidia and ascospore measurements were recorded and compared across the three hosts. Because macroconidia were absent in strains originating from tulip-poplar, no formal measurements were taken for any of the strains used in this study even when present (SUPPLEMENTARY FIG. 2).

Field collected bark plugs with perithecia were selected based on ascocarp maturity and density. Ascospores averaged 13.2 μm (10.2-15.66 μm) in length across the 6 samples examined (FIG. 1C,G,K, FIG. 6A). Among these measurements, mean spore length was significantly larger for *N. magnoliae* ascospores originating from tulip-poplar (13.8 μm) compared to *N. magnoliae* ascospores originating from either Fraser magnolia (13.2 μm) or star magnolia (12.6 μm) based on a 1–way ANOVA (*F*_2,147_ = 20.143, *P* < 0.001) and subsequent Tukey’s HSD test with Bonferroni correction (FIG. 6A). Ascospore length was smallest from star magnolia.

Microconidia averaged 7.5 μm (4.67-13.43 μm) in length and 2.8 μm (1.97-3.74 μm) in width across the 9 isolates examined (FIG. 6B-C). Among these isolates, significant differences in microconidia length were found between at least 2 groups of strains based on Welch’s ANOVA (*F*_2,281_ = 23.5, *P* < 0.001). The Games-Howell post-hoc test revealed significant differences in microconidia length between tulip-poplar (7.1 μm) and Fraser magnolia (7.9 μm) strains (adjusted *p*-value = < 0.001) and significant differences between Fraser magnolia (7.9 μm) and star magnolia (7.1 μm) strains (adjusted *p*-value = < 0.001) (FIG. 6B). However, no significant differences in microconidia length were observed between tulip-poplar and star magnolia strains. Additionally, we found that there was a significant difference in microconidia width between at least 2 groups of strains after conducting a 1-way ANOVA (*F*_2,447_ = 27.21, *P* < 0.001). The Tukey’s HSD post hoc test revealed significant differences in microconidia width between tulip-poplar (2.7 μm) and Fraser magnolia (2.9 μm) strains and significant differences between tulip-poplar (2.7 μm) and star magnolia (3.0 μm) strains (FIG. 6C). Furthermore, it was observed that our singular star magnolia strain of *N. magnoliae* also abundantly produces macroconidia similar to Fraser magnolia strains, though measurements and average number of septations were not recorded (SUPPLEMENTARY FIG. 2).

**Figure 6.**
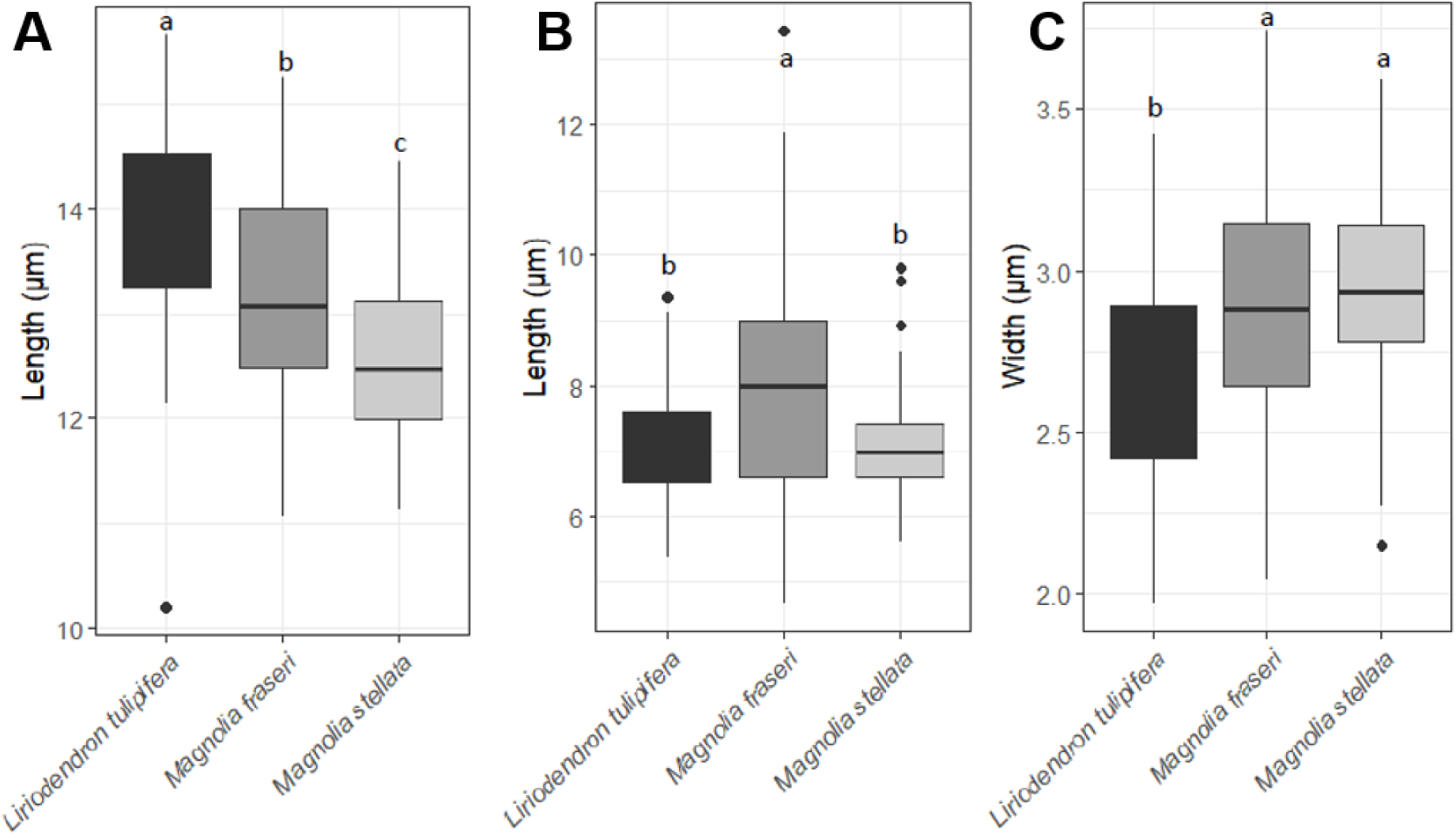
Boxplots of ascospore length (A) and microconidia length (B) and width (C) for strains of *Neonectria magnoliae.* Strains are grouped by host origin (X-axes). Letters above plots represent significant.

#### Taxonomy

*1) Neonectria magnoliae* f. sp. *liriodendri* H.M. Petronek, M.T. Kasson, C.M. Stauder, f. sp. nov., holotype info: Mycobank #288753

*Material Examined:* Single ascospore PDA cultures originating from fresh perithecia from cankered *Liriodendron tulipifera* (tulip-poplar) trees at Fernow Experimental Forest, Tucker County, West Virginia, USA in July 2018 (**NRRL 64651**) and from Snake Hill Wildlife Management Area, Monongalia County, West Virginia, USA in July 2018 (**NRRL 64649**), collected by C.M. Stauder and M.T. Kasson. Additional strains examined include NmLtBg_02 and NmLtBg_04 from West Virginia Botanic Garden in Monongalia County and NmLtAu_01 from Audra State Park in Barbour County.

*Note:* Mycobank number references the holotype strain CBS 380.50 / BPI 552527 / No. 64187 recovered from cankered *L. tulipifera* in Bent Creek Experimental Forest in Asheville, NC on October 12, 1934 by M. L. Lohman. No pathogenicity / virulence tests have been formally conducted on the holotype strain but is included as *N. magnoliae* f. sp. *liriodendri* based on host origin and 100% sequence similarity in the nuclear ribosomal RNA gene repeat (rDNA) comprising the internal transcribed spacer region (*ITS:* **MH856671**) and the RNA polymerase II second largest subunit (*RPB2:* **HQ897713**) gene sequences for CBS 380.50 and NRRL 64651. GenBank accession numbers associated with strains from this study are as follows: TBD.

*Etymology:* Named after the host (*Liriodendron tulipifera*, botanical name) from which it was isolated.

*2) Neonectria magnoliae* f. sp. *magnoliae* H.M. Petronek, M.T. Kasson, C.M. Stauder, f. sp. nov.

*Material Examined:* Single ascospore cultures originating from fresh perithecia from cankered Fraser magnolia (*Magnolia fraseri*) trees at Gaudineer Knob Scenic Area, Monongahela National Forest, Randolph County, West Virginia, USA in July 2020 (NRRL 64650 **/** GKMAG7202) and GKMAG5 in July 2020, collected by C.M. Stauder and M.T. Kasson and from West Virginia University’s Core Arboretum in Monongalia County, West Virginia, USA in September 2023 (NmMfCa_01). GenBank accession numbers associated with strains from this study are as follows: TBD.

*Etymology:* Named after the host (*Magnolia fraseri,* botanical name) from which it was isolated.

## DISCUSSION

In this study, we sampled and characterized the diversity of *N. magnoliae* strains across three hosts spanning seven counties in West Virginia. The results of this work unocvered two subspecific pathotypes in *N. magnoliae*: *Neonectria magnoliae* f. sp. *liriodendri* and *Neonectria magnoliae* f. sp. *magnoliae*. *Neonectria magnoliae* f. sp. *liriodendri* occurs exclusively on tulip-poplar with increased virulence on this host, while *Neonectria magnoliae* f. sp. *magnoliae* occurs exclusively on *Magnolia* spp. with increased virulence on this Fraser magnolia.

Lohman & Watson (1943) previously reported *N. magnoliae* from tulip-poplar in Connecticut, Ohio, North Carolina, Tennessee, Virginia, and West Virginia, from Fraser magnolia in Tennessee and West Virginia, and from umbrella magnolia in West Virginia. Except for *N. magnoliae* holotype strain CBS 380.50 from tulip-poplar in North Carolina, nucleotide sequence data is not publicly available for the paratype from Richwood, WV, the other paratypes, or any contemporary isolates outside West Virginia. Sequence data for a single *N. ditissima* strain recovered from tulip-poplar in Tennessee indicates that *N. ditissima* overlaps with at least part of the historic range for *N. magnoliae* (Lohman and Watson 1943). Other red perithecia-producing members of the Nectriaceae have also been reported from tulip-poplar including *Neocosmospora silvicola* from Tennessee and *Ilyonectria liriodendri* from California, though the latter falls outside the reported historic range of *N. magnoliae* (Chaverri et al. 2011, Sandoval-Denis et al. 2019). Nevertheless, the co-occurrence of other phytopathogenic genera and species with overlapping ascocarp features undoubtedly contribute to confusion regarding the geographic range of these species and whether the range of *N. magnoliae* on tulip-poplar is more restricted than the literature indicates.

Both methods of phylogenetic inference for both the single-locus trees and concatenated trees resolved *N. magnoliae* as a strongly supported monophyletic group. The single-locus phylogeny for *EF1-α* provided additional intraspecies resolution for *N. magnoliae*, revealing a single moderately supported clade containing all tulip-poplar isolates (SUPPLEMENTARY FIG. 1C). However, this topology was incongruent with all other individual loci and both concatenated datasets (SUPPLEMENTARY FIG. 1A-B,D). The failure to generate *EF1-α* sequences for three critical magnolia-derived strains despite repeated attempts contributed to this uncertainty. Interestingly, Fraser magnolia isolates, though limited in number, show considerable sequence divergence based on geography, a pattern which is not seen in isolates originating from tulip-poplar despite the inclusion of more isolates with broader geographic distance. This possibly may be due to a discontinuous host range which might limit gene flow between geographically isolated populations. Given the incongruence among single-gene phylogenies, these data fail to meet the criteria for the Genealogical Phylogenetic Species Recognition principle (GCPSR) (Taylor et al. 2000). As such, our current data do not support *N. magnoliae* as two species but instead support host-specific pathotypes that specialize on either tulip-poplar or Fraser magnolia.

Follow-up studies that include additional loci (e.g. *RPB1, RPB2, GDP*) are needed. Towards this end, we recently generated genomic resources for West Virginia strains of *N. magnoliae* from tulip-poplar (NRRL 64651) and *N. punicea* from sugar maple (NRRL 64653) to allow for in-depth comparisons (Petronek et al. 2024). Ongoing efforts are focused on generating additional genomic resources for Fraser magnolia strains of *N. magnoliae.* Despite the economic impacts of *Neonectria* fungi and closely allied species, much remains unknown about the genetic diversity and distribution of these nectriaceous canker pathogens (Ghasemkhani et al. 2016).

The results of our tulip-poplar comparative pathogenicity and virulence assays were not consistent with the findings of Spaulding and colleagues (1936) on this same host: significant differences in canker sizes were not noted between isolates originating from either host. Interestingly, Fraser magnolia strains produced larger cankers on Fraser magnolia than tulip-poplar strains and likewise, tulip-poplar strains produced larger cankers on tulip-poplar than Fraser magnolia strains. Given the differences in virulence observed on their respective hosts, we propose two subspecific pathotype designations for *N. magnoliae*: *Neonectria magnoliae* f. sp. *liriodendri* and *Neonectria magnoliae* f. sp. *magnoliae*.

Though these observations certainly support host-associated preferences, the possibility of a host jump by *N. magnoliae* from Fraser magnolia to tulip-poplar remains unclear. Nevertheless, several lines of evidence generally support this hypothesis. Strains originating from tulip-poplar lack macroconidia production, have less sequence variation despite broader sampling, and appear to cause increased mortality of tulip-poplar, especially small diameter trees with intermediate or overtopped crown classes.

In this study, we uncovered a previously unreported host for *N. magnoliae,* star magnolia (*Magnolia stellata*), an ornamental shrub planted widely throughout the region. This is to our knowledge the first report of this native pathogen on a non-native Magnoliaceae host. Notably, this non-native host was planted in the same vicinity as a Fraser magnolia in WVU’s Core Arboretum, which also had branch cankers caused by *N. magnoliae*. Despite their proximity and shared magnolia hosts, these strains were not genetically similar compared to other strains recovered together. For example, comparisons among multiple strains from multiple cankered Fraser magnolia trees sampled at Gaudineer Knob were nearly identical based on the loci examined. The progression of disease and associated dieback on this non-native host seems much more significant compared to the two native hosts. Pathogenicity trials on star magnolia are needed, but as non-native ornamentals, locations with sufficient numbers of plants for field studies are rare. Conversely, magnolias can be purchased from nurseries for experiments, but can be prohibitively expensive if numerous plants are needed. Other magnolia hosts should also be surveyed for these nectriaceous canker pathogens, like umbrella magnolia (*Magnolia tripetala*) and southern magnolia (*Magnolia grandiflora*), given recent reports of cankers of unknown origin on these hosts (Pettis 2022).

Historically, ascospore measurements have been used for characterizing species within *Neonectria* and closely related genera (Lohman & Watson 1943; Booth 1959; Kasson & Livingston 2009). In this study, our ascospore & microconidia measurements for *N. magnoliae* were in agreement with those provided by Lohman & Watson (1943) for the holotype and paratype specimens. Within *N. magnoliae*, strains from tulip-poplar had significantly larger ascospores compared to strains from Fraser and star magnolia. Ascospores from strains originating on Fraser magnolia also had significantly larger ascospores compared to those strains originating on star magnolia. While average ascospore length distinguishes *N. magnoliae* from other *Neonectria* species, these measurements are still variable, making it difficult to diagnose the cause of a ‘Nectria’ canker based on ascospore measurements alone. For example, researchers reported *Neonectria* mean ascospore measurements from American beech samples in northern Maine that overlapped with measurements recently reported for *N. magnoliae* ascospores originating from perithecia from tulip-poplar (Kasson & Livingston 2009). Given the lack of magnolia hosts in norther Maine, these findings do not support the presence of *N. magnoliae* f. sp. *liriodendri*. However, these findings together demonstrate the limitations of fungal identification based exclusively on select morphological features such as ascospore length.

While conidia measurements have not historically been used for identification, the abundance of microconidia from all isolates allowed for meaningful comparisons. Microconidia were significantly larger (in both length and width) for Fraser magnolia isolates compared to those from tulip-poplar hosts. The star magnolia isolate was intermediate, with similar length as tulip-poplar isolates and similar width as Fraser magnolia isolates. Macroconidia were only produced by strains originating from Fraser magnolia or star magnolia. Since this study focused on comparisons between the two primary hosts, macroconidia measurements were not taken given the lack of macroconidia in tulip-poplar strains. It is surprising that despite wide geographic sampling in West Virginia, tulip-poplar strains always lack macroconidia, even after months of growth. Another *Neonectria* species from China, *N. microconidia,* is also reported to exclusively produce microconidia in culture (Zhao et al. 2011). Future studies should investigate if the loss of the macroconidia spore stage is disadvantageous to the fungus, and how the loss impacts the disease cycle of these species.

Morphological and phylogenetic data were limited from star magnolia as only one symptomatic tree was found despite additional surveys on this host in the greater Morgantown area. As such, we were unable to resolve its relationship among strains from the other two hosts. However, the production of macroconidia supports its *Magnolia* host-origin. As star magnolia is a common ornamental shrub planted in suburban landscapes, opportunities abound to survey this host for decline and branch and stem cankers.

Through comprehensive studies, two morphologically distinct formae speciales in *Neonectria magnoliae* are described that differ in virulence between two Magnoliaceae hosts. Multi-locus phylogenetic studies found insufficient evidence for two distinct species within *N. magnoliae*. However, both formae speciales in *N. magnoliae* appear adapted to their respective hosts, based on comparative pathogenicity and virulence testing. The formae speciales are also morphologically distinct, even in geographic regions where both hosts overlap.

## DATA AVAILABILITY STATEMENT

The data that supports the findings of this study are available in the supplementary material of this article or are openly available as sequence data deposited as Sanger sequences (Accessions TBD) in National Center for Biotechnology Information. All data and code for analyses included in this manuscript are available on GitHub: https://github.com/(To Be Deposited).

## ACKNOWLEDGEMENTS

We thank Hana Barrett, Colin Gibney, Andrew Kasson, Amy Metheny, Kristen Pierce, Molly Sherlock, and Cameron Wilson for their assistance with help with field inoculations. We thank Dr. Jamie Schuler and Forest Manager Heidi Harmala for helping coordinate use of WVU’s University Forest for field pathogenicity experiments.

## SUPPLEMENTARY FIGURES

**Supplementary Figure 1.**
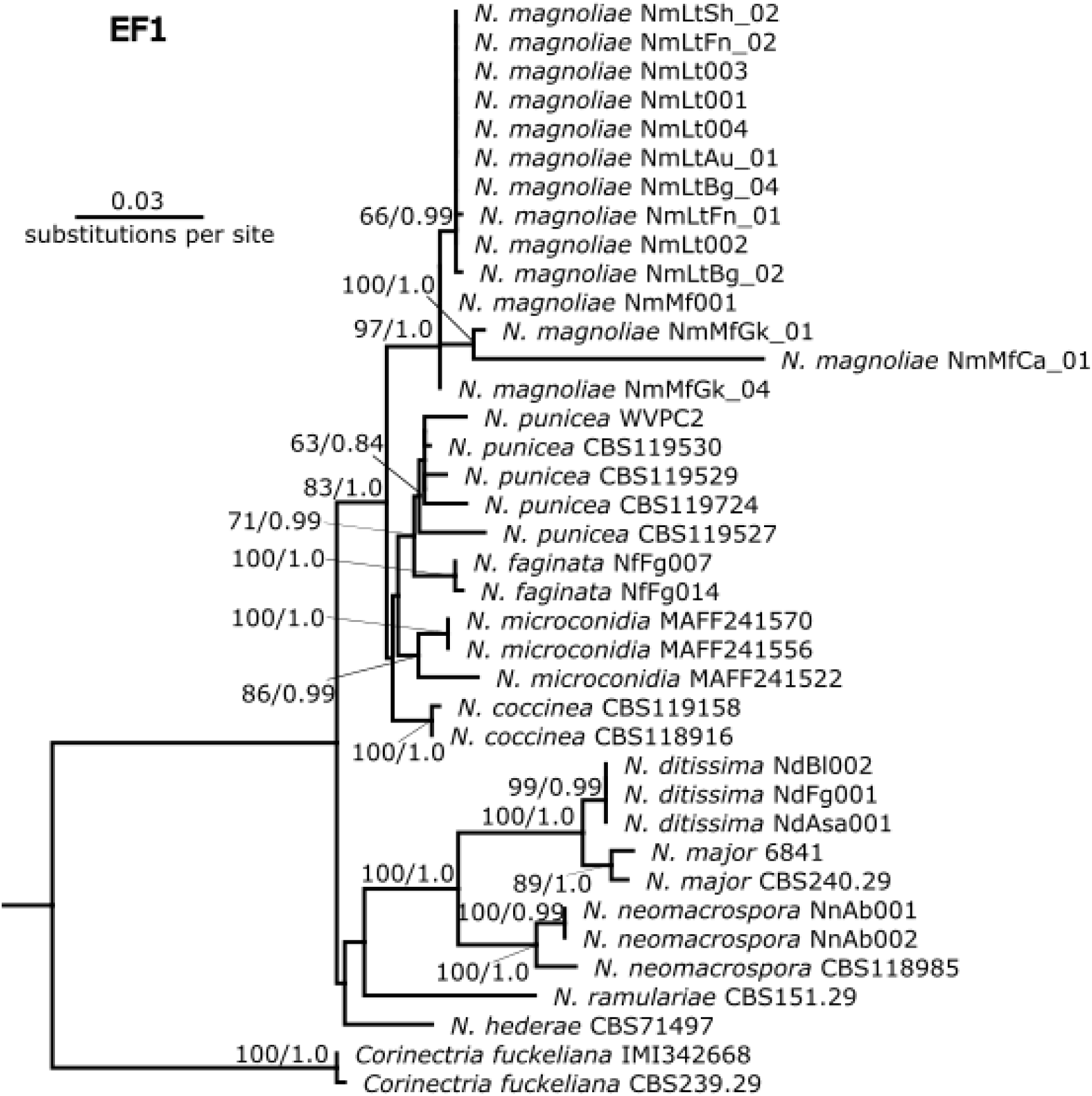
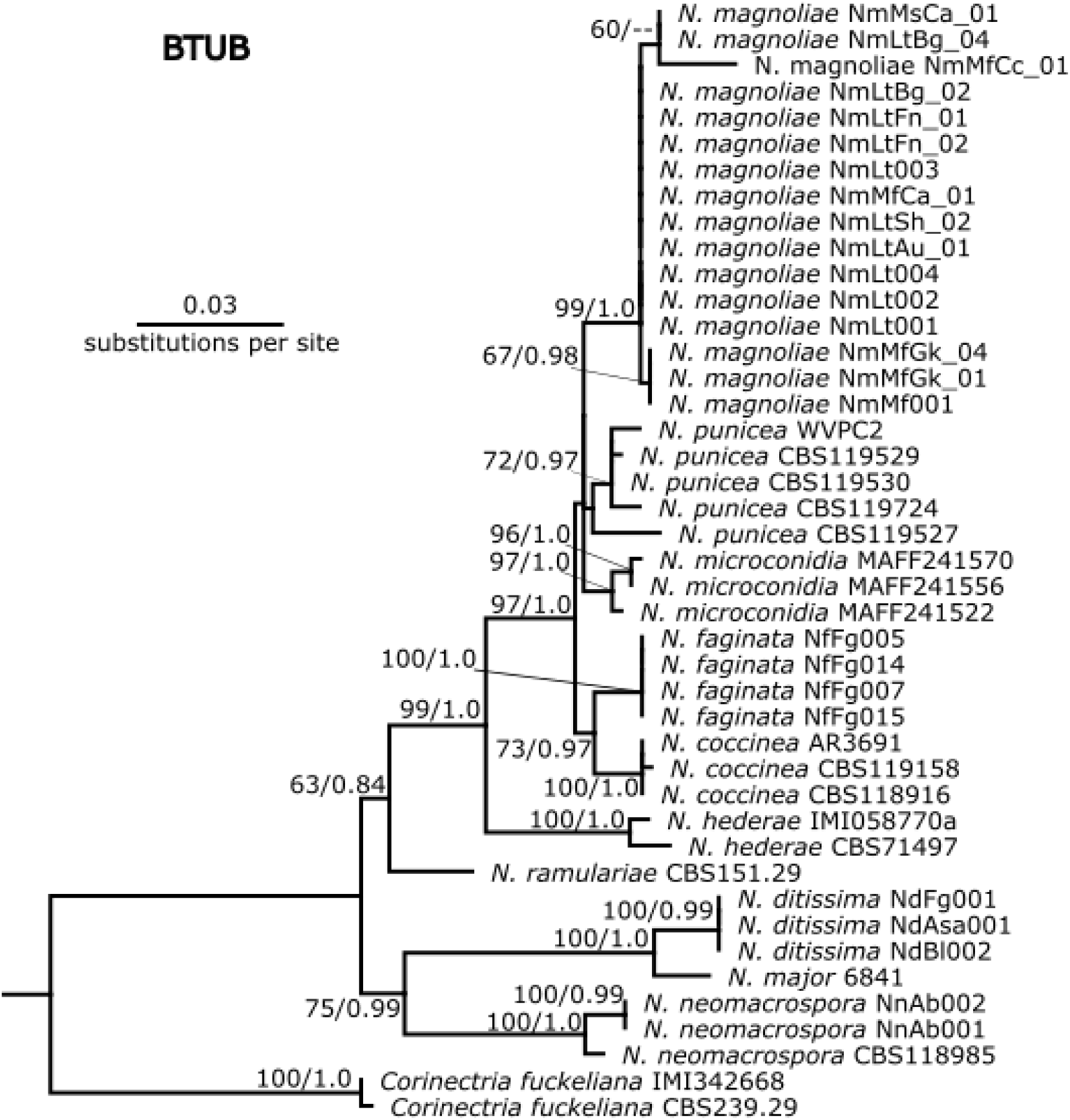
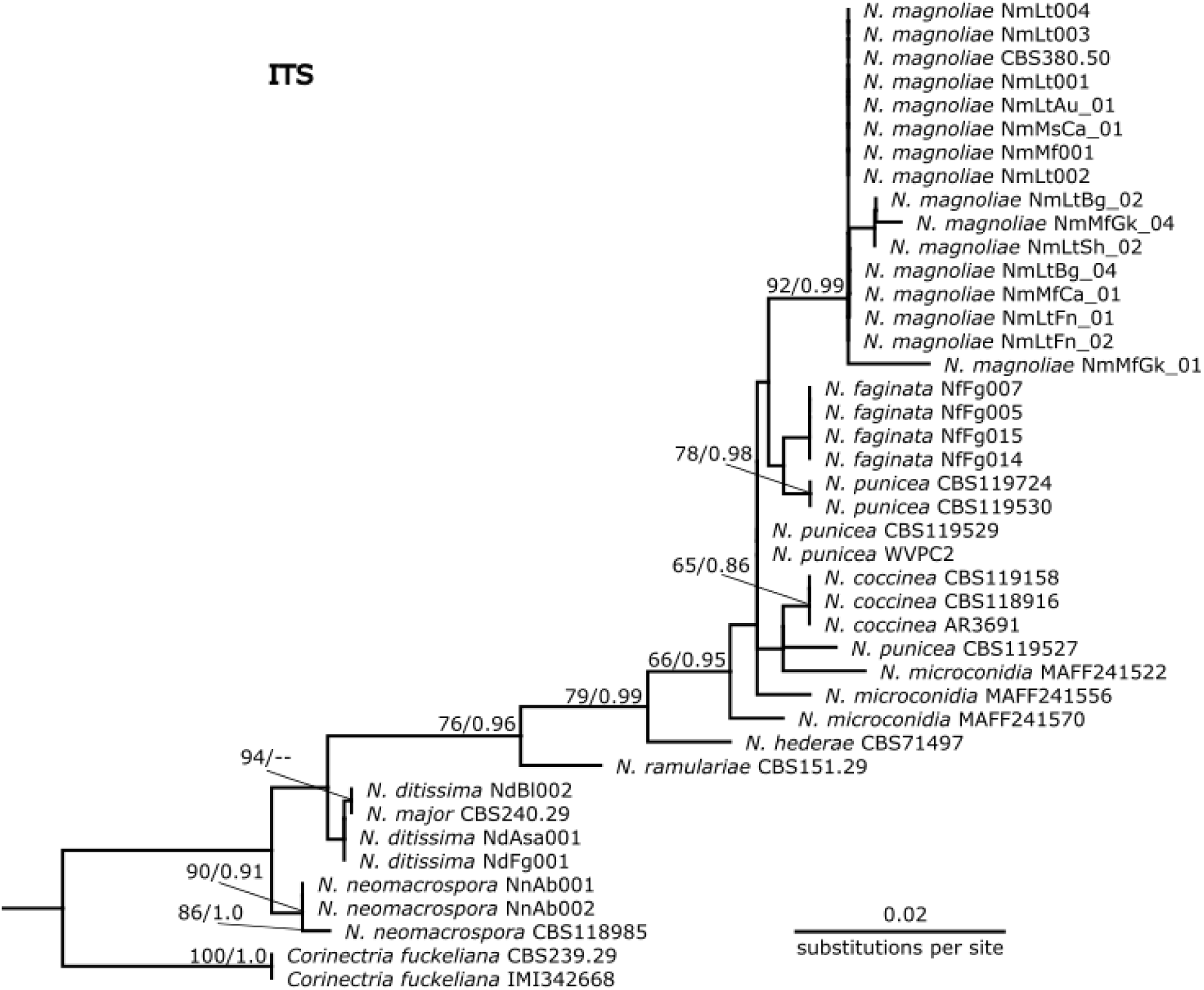
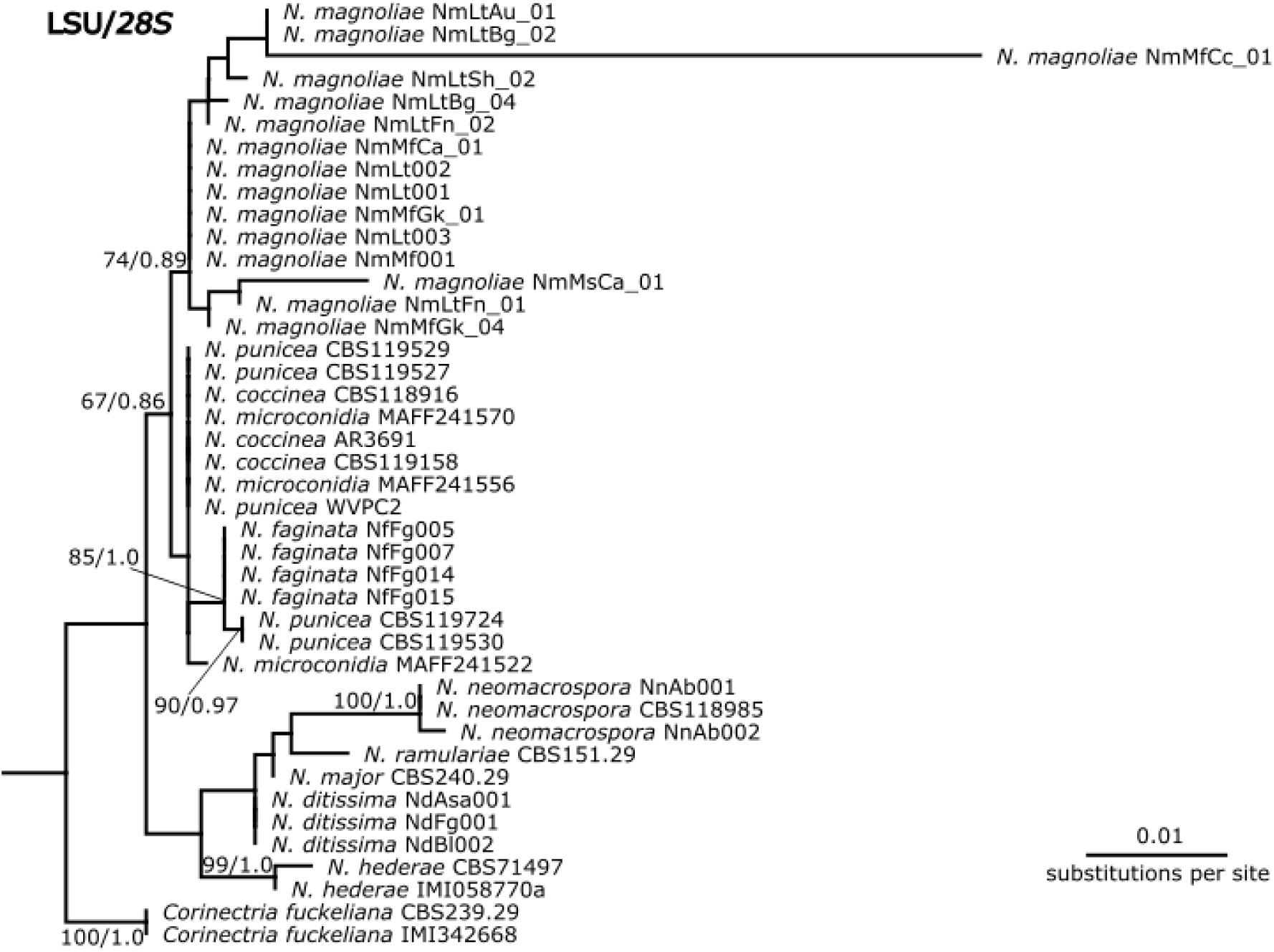
Single-gene phylogenetic trees for *EF1-α* (A)*, BTUB* (B)*, ITS* (C), and *28S* (D) for *Neonectria* spp. with *Corinectria fuckeliana* as the outgroup. Topology and branch lengths are from the ML analysis. For each node supported in the ML analysis, bootstrap support and posterior probabilities are indicated (ML/BI) with one exception. Bootstrap support values < 50 for ML analysis are not shown nor are their corresponding BI posterior probability values. Dashes indicate that the node did not appear in the BI analysis. Strain metadata including GenBank accession numbers are listed in TABLE 1.

**Supplementary Figure 2.**
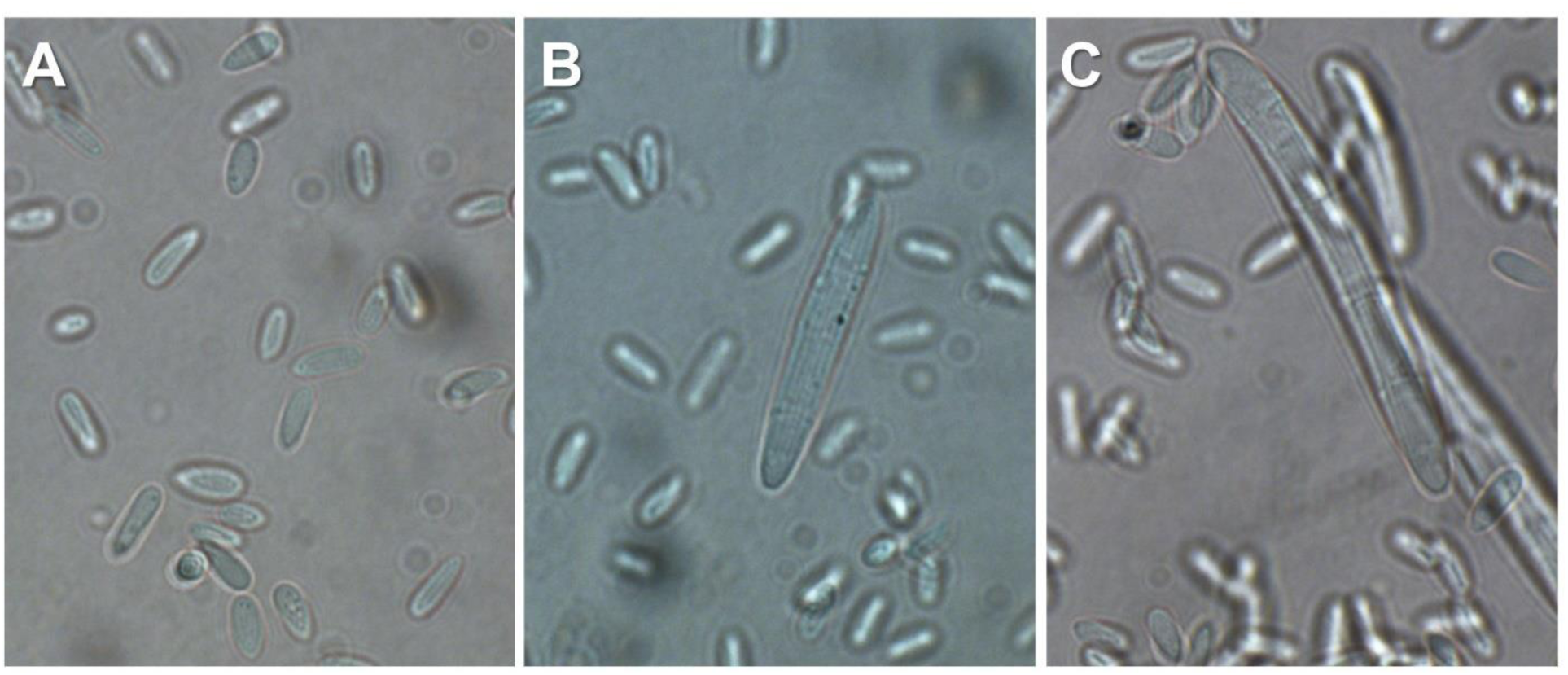
Microconidia of *N. magnoliae* strains from a tulip-poplar (NRRL 64651; A), Fraser magnolia (NRRL 64650; B) and star magnolia (NRRL 64648; C). Macroconidia (large multi-septate spores in B-C) were only observed in *Magnolia* strains. All microscopy images were taken at 100x oil magnification.

